# Origins of major archaeal clades do not correspond to gene acquisitions from bacteria

**DOI:** 10.1101/019851

**Authors:** Mathieu Groussin, Bastien Boussau, Gergely Szöllősi, Laura Eme, Manolo Gouy, Céline Brochier-Armanet, Vincent Daubin

## Abstract

In a recent articxle, Nelson-Sathi et al. [NS] report that the origins of Major Archaeal Lineages [MAL] correspond to massive group-specific gene acquisitions via horizontal gene transfer (HGT) from bacteria (Nelson-Sathi et al., 2015, Nature 517(7532):77-80). If correct, this would have fundamental implications for the process of diversification in microbes. However, a re-examination of these data and results shows that the methodology used by NS systematically inflates the number of genes acquired at the root of each MAL, and incorrectly assumes bacterial origins for these genes. A re-analysis of their data with appropriate phylogenetic models accounting for the dynamics of gene gain and loss between lineages supports the continuous acquisition of genes over long periods in the evolution of Archaea.

## Results and Discussion

Reconstructing genome histories is a major challenge in evolutionary biology and the subject of a large body of literature (Maddison 1997; Snel et al. 2002; Mirkin et al. 2003; Hahn 2007; Csűrös 2010; Boussau et al. 2013; Szöllõsi et al. 2013). In a recent study, NS devised an *ad-hoc* method to infer ancestral gene acquisitions in the history of Archaea (Nelson-Sathi et al. 2015). From 134 archaeal proteomes, they built 25,762 protein families of which 2,264 had at least two representatives in a single MAL (forming a monophyletic group) and at least two bacterial homologs (out of 1,847 genomes) belonging to species from two different phyla. NS concluded that all 2,264 gene families were acquired from Bacteria at the origin of MALs, implying that these acquisitions probably promoted their origin and evolutionary success.

A close look at the results of NS is enough to convince oneself that there are problems with their approach. The set of genes that NS infer to have been acquired at the roots of MAL comprises 2,264 gene “clusters”, which are called “import” clusters. Fig. 1a presents the tree reconstructed from one of them, “Cluster 23981”, which we simply sampled from the list of “import” clusters available as supplementary material accompanying NS (table 3 from supplementary data) (Nelson-Sathi et al. 2015). This gene is found in only two sister species from a single archaeal genus (*Methanosarcina*), nested in the order Methanosarcinales (one of the 12 MALs) (Petitjean et al. 2015), and two very distantly related Bacteria (*Bradyrhizobium japonicum*, an alphaproteobacterium and *Granulicella tundricola*, an acidobacterium), from two different phyla (out of 1,981 genomes in total). Inclusion of this cluster in the “import” set implies that, according to NS: i) the gene was transferred before the origin of Methanosarcinales and ii) that it is “very widespread among diverse bacteria, clearly indicating that [it is an] archaeal acquisition from bacteria, or import” (Fig. 1) (Nelson-Sathi et al. 2015). The first is akin to saying that a gene present only in chimpanzee and human necessarily originated in the ancestor of all Vertebrates and has since undergone *systematic* gene loss in *all* other vertebrate species. Similarly, a “widespread” distribution (ii) signifies transfer from a bacterial donor only if it is interpreted as a sign of antiquity in Bacteria, again implying extensive gene losses, this time in Bacteria. A much more parsimonious scenario (Fig. 1) requires only two transfers, and avoids massive convergent losses in Archaea and Bacteria. In this scenario, the gene is acquired at the origin of the genus *Methanosarcina* (but not the phylum Methanosarcinales), and the direction of HGT is unknown.

**Figure 1:**
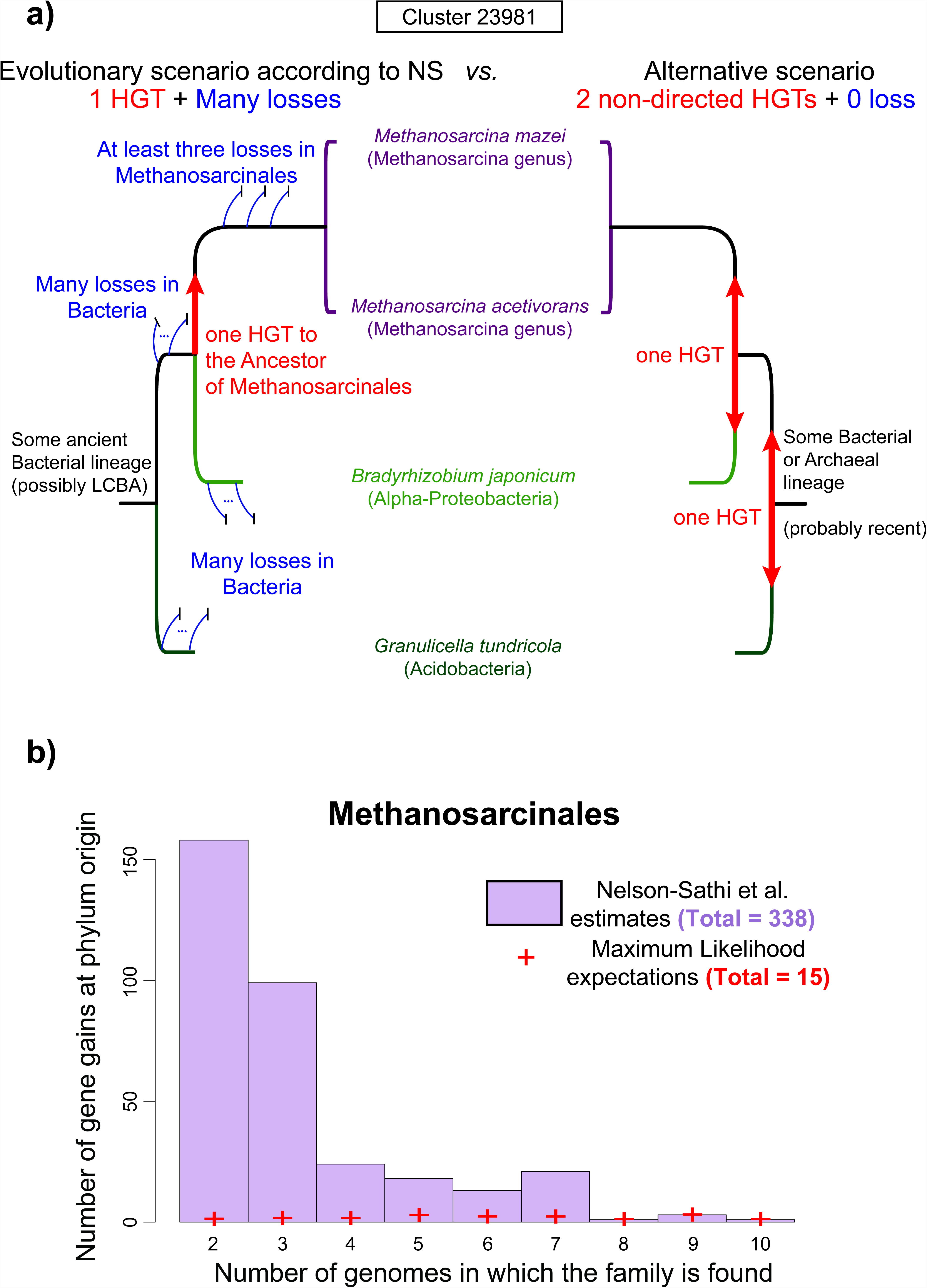
Shoehorning HGTs to the origins of archaeal phyla. a) Competing evolutionary scenarios for « Cluster 23981 » from NS’s ‘import’ set. Left: the gene is ancient in Bacteria and was subsequently transferred (red arrow) to the ancestor of Methanosarcinales. A large number of losses (blue lines) in Bacteria and Methanosarcinales is necessary to explain the narrow pattern of presence in extant species. Right: the gene is not ancient in Bacteria and was absent at the origin of Methanosarcinales. It has been transferred twice from Bacteria or Archaea and among Bacteria, and no loss is necessary. b) Distribution of gene gains at the origin of Methanosarcinales (for other phyla, see the Appendix). NS estimates are represented in purple. The distribution is very skewed towards sparsely distributed genes. Maximum Likelihood expectations of gains on the corresponding set of genes are represented by red crosses.

Unfortunately, this cluster is not an exception but is representative of the “import” set (Fig. 1b, Fig. 2 and Supp Fig. 1). Most of the genes reported by NS as acquired at the origins of a MAL are present in very few species in Archaea and Bacteria. In fact, 52% (1,171/2,264 “import” clusters) are represented in only two or three archaeal species, strongly suggesting that these genes have been acquired during the diversification of MALs rather than at their root (see below). Furthermore, the definition of “import” genes by NS requires that they have homologs in bacterial species from at least two phyla, of which they claim one has to be the donor (Nelson-Sathi et al. 2015). Although this is not explicit in their paper, it can only mean, as we argue above, that NS consider that if a gene has representatives in two different bacterial phyla, it is *ancient* (*i.e.*, it was present in the common ancestor of these two phyla) and hence of bacterial origin. Yet, these “import” genes could be more recent and instead could have been transferred among Bacterial phyla. In support of this hypothesis, we observe that these genes have a very narrow and patchy distribution: half of these “import” genes have homologs in less than 1.1% of the bacterial genomes considered (21/1847) (Fig. 2). Because these genomes are from at least two phyla, such a patchy distribution is consistent with, and strongly suggests, recent HGTs within the bacterial domain. Their presence in a common ancestral bacterial genome cannot be assumed. Instead, these genes appear to have very complex evolutionary histories, and NS’s assumption that they were transferred from Bacteria to Archaea rather than the reverse is unfounded.

How did NS come to their conclusions about the origins of these genes? To assess whether the 2,264 gene acquisitions correspond to the origins of MAL, they employed an *ad hoc* phylogenetic test, which compares distributions of splits in the “import” and “recipient” set of gene trees. The “recipient” set is comprised of gene families only present in a single MAL, whereas members of the “import” set, discussed above, also have homologs in Bacterial species. NS show that the “import” and “recipient” sets exhibit similar distributions of splits for 6 out of 13 MALs. They interpret this result as evidence that the “import” set of genes has been vertically inherited after a single acquisition at the root of the corresponding MAL. In reality, this result only shows that tree distributions are not statistically different between these two — arbitrary — sets of genes. This similarity *does* imply that gene transfer rates are similar between the two sets. It *does not* imply that either set was predominantly acquired above the root of a particular MAL. This would only be the case if genes of one of the sets (e.g. the “recipient” set) were predominantly acquired above the root of the given MAL. In reality, both “recipient” and “import” genes have a very skewed distribution: respectively 59% and 52% of these genes are present in less than four species of the MAL. Furthermore, this test is only applicable to families with four or more genes (see the Supplementary Methods of NS), which only represent 48% of the 2,264 families in the “import” set. Nevertheless, NS extend their conclusions to *all* genes in the import set. Finally, while 7 out of 13 MALs do not pass their congruence test for a lack of statistical signal, NS nonetheless argue that *all* 2,264 import genes were acquired at the origins of 13 MALs (as clearly stated, for instance in the abstract or the caption of Fig.3 in Nelson-Sathi et al. (2015)).

**Figure 2:**
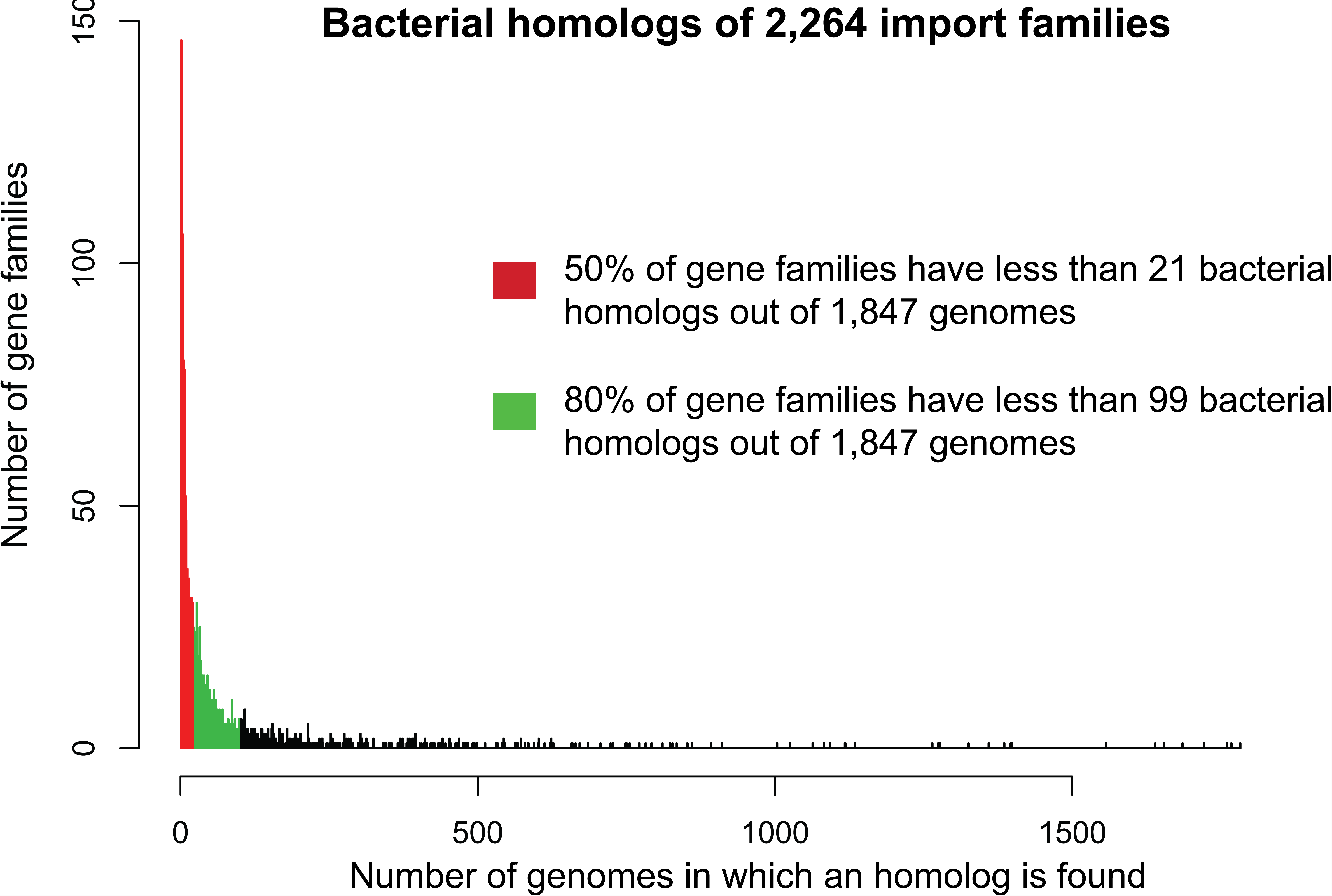
The sporadic distribution of the 2,264 “import” gene families in Bacteria. The distribution of the 2,264 gene families in the 1,847 bacterial genomes is represented. The distribution is very skewed: half of the gene families have fewer than 21 bacterial homologs out of the 1,847 genomes (1.1%), and a vast majority of them (80%) are present in fewer than 99 bacterial genomes (5%). Because for each family the bacterial homologs are from at least two different phyla, this distribution is highly suggestive of recent HGTs among bacteria and of complex evolutionary scenarios for these families, preventing the inference of a direction of transfer.

**Figure 3:**
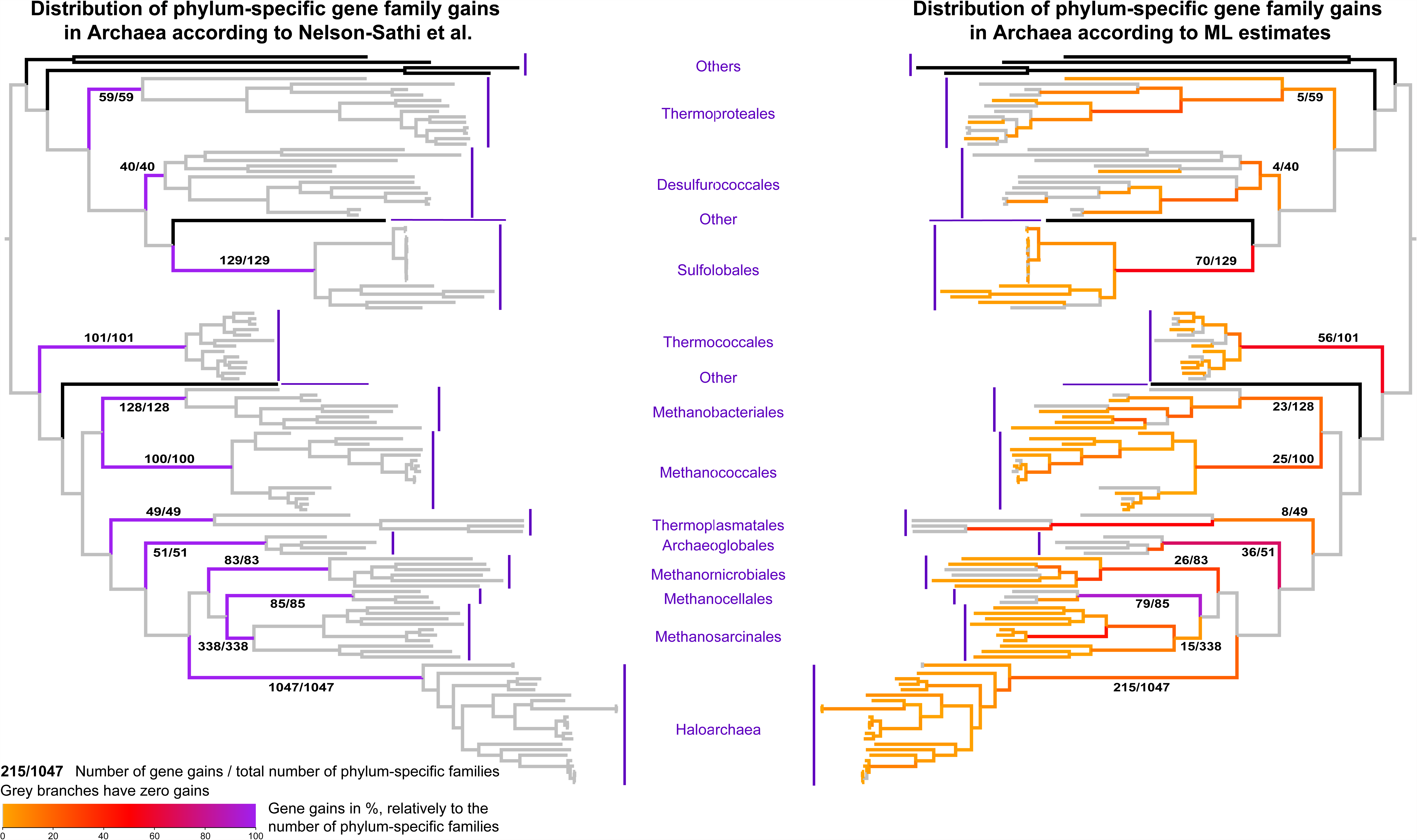
Most of the 2,264 “import” gene families were acquired after phyla origination in Archaea. We used the tree reconstructed by NS to represent the points of acquisition of genes of the “import” set in the evolution of Archaea. Colors on branches represent the expected number of gains per branch, relatively to the number of phylum-specific families. Branches with a gain expectation < 1 are colored in grey. On the left, NS estimations are represented. All phylum-specific families were acquired at the origin of each phylum, and no subsequent gains are inferred. On the right, Maximum Likelihood (ML) estimates show that gene family gains are spread over the history of diversification of each phyla and that most of the families were acquired after the origination of phyla

In fact, with the method used by NS, no gene acquisition is possible after the ancestor of a MAL because the relationships among species within MALs are ignored. In order to assess how many of the genes specific to MALs have been acquired more recently, it is necessary to analyze their data in a phylogenetic framework. Ideally, it would be necessary to apply a method that simultaneously infers the species tree and the scenarios of gene evolution for each cluster based on the corresponding gene trees (Szollosi et al., 2012). However, these methods require extensive computation and are currently limited in the number of species that they can efficiently analyze. We hence used Count (Csűrös 2010), using a maximum likelihood approach with an evolutionary model of gene gain and loss (see Methods) on the phylogenetic profiles of each gene cluster. We estimated lineage-specific rates of gain per gene cluster and computed the expected lineage-specific number of gains along the reference tree reconstructed by Nelson-Sathi et al. (2015). This approach is of course a simplification, because we assume that the tree of species within MALs is correct and we ignore the phylogenetic signal contained in individual gene alignments. Yet, it is conservative for our comparison here because errors in the relationships of species within a MAL and ignoring the phylogenetic conflict between a gene tree and the species tree will both tend to yield a higher probability for genes to be present at the origins of MALs. Among the 2,264 “import” genes, the great majority (75%) appear *during* the diversification of each MAL (Fig. 3 and Supp. Fig. 1). In other words, the acquisitions of the “import” genes defined by NS are spread over long periods in the evolution of MAL, and only a minority (25%) can be inferred to have been acquired at their origins. For instance, the number of gene acquisitions at the root of Methanosarcinales and Thermoproteales is now very small (15 instead of 338 and 5 instead of 59, respectively). Finally, in most MALs (Haloarchaea, Methanosarcinales, Methanomicrobiales, Thermoplasmatales, Methanobacteriales, Desulfurococcales and Thermoproteales), some internal branches experienced significantly more gains than the branch leading to their last common ancestor (Fig. 3). These estimates are consistent using ML or parsimony with a variety of parameter values (see Methods and Supp Fig. 2).

The approach used by NS was first applied in a previous study that found that massive gene transfers had occurred at the origin of Haloarchaea (Nelson-Sathi et al. 2012). These conclusions have since been shown to be unfounded (Becker et al. 2014). Gene transfer seems to have occurred continuously across the tree of life (Ochman et al. 2000; Gogarten and Townsend 2005; Abby et al. 2012; Szöllõsi et al. 2012), and probably contributed to the greatest as well as the smallest innovations. Understanding the evolutionary history of these adaptations requires and deserves more accurate analysis of the data in an integrated phylogenetic framework.

## Methods

The data used in Nelson-Sathi et al. (2015) were kindly provided by the authors. Statistical analyses were performed in R (R Core Team, 2013) and ancestral gene repertoire reconstructions were performed with the Count program (Csűrös 2010). Branch-specific rates of a birth/death model of gain and loss of gene clusters (Csűrös and Miklós 2006) were estimated using Maximum Likelihood from the 25,762 protein families, along the archaeal species tree reconstructed by Nelson-Sathi et al. (2015). After optimization of the branch-specific parameters, ancestral reconstructions were carried out for each of the 2,264 families, by computing branch-specific posterior probabilities of evolutionary events. At a given branch, the expected number of gene acquisitions was computed by summing all family-specific gene gain posterior probabilities. Wagner parsimony (Farris 1970; Csűrös 2008) with a large range of gain and loss cost combinations was also tested. All parsimony and ML estimations gave very similar results (See Supp Fig. 2). A Count session file that will allow users to reproduce our results is available at ftp://pbil.univlyon1.fr/pub/datasets/DAUBIN/HGT_Archaea/.

